# Discovery of *Bordetella* oligosaccharide – extracellular glycan common to genus *bordetella*. Structure, immunogenicity and possible implications for host-pathogen interactions

**DOI:** 10.64898/2026.04.29.721555

**Authors:** Karolina Ucieklak, Sabina Koj, Tomasz Niedziela

**Author notes:** Corresponding author: Tomasz Niedziela; Department of Immunochemistry, L. Hirszfeld Institute of Immunology and Experimental Therapy, Polish Academy of Sciences, R. Weigla 12, PL-53-114 Wroclaw.

## Abstract

Pertussis pathogenesis is the result of multiple virulence factors. In addition to the secretory proteins of *Bordetella pertussis*, surface molecules such as adhesins and endotoxin play a role in the pathogenesis of this disease. There are conflicting reports on the existence and nature of the *Bordetella* capsular polysaccharides or exoglycans. The data concerning the glycome of *Bordetellae* is incomplete. This conclusion is primarily derived from genomic data, with only limited indications regarding the actual structures. In this study, we present novel data on the exoglycan produced by all strains and species of the investigated bacteria from the genus *Bordetella*, including *Bordetella pertussis, Bordetella parapertussis, Bordetella bronchiseptica,* and *Bordetella holmesii*. This is the first time this type of data has been provided. The exoglycan was consistently recovered from the chemically defined culture media of various *Bordetellae* species and strains. The compound was identified by nuclear magnetic resonance (NMR) as a free hexasaccharide released into the medium and thus received its name, *Bordetella* oligosaccharide (BOS). The biosynthetic origin of the BOS was confirmed by NMR combined with metabolic labeling in culture, using ^13^C,^15^N-L-glutamate as a primary carbon source. The identification of BOS has the potential to enhance our comprehension of the complete array of virulence factors contributing to the pathogenesis of *Bordetella pertussis*, particularly in regard to their relations with other *Bordetella* species. In the field of vaccine design, glycan structures are typically of utmost importance; however, they were hardly ever considered in the case of pertussis.

## Introduction

The genus *Bordetella* comprises a group of aerobic Gram-negative small coccobacilli. Most of these bacteria have adapted to live in a close relation with higher organisms. *B. pertussis*, *B. parapertussis, B. bronchiseptica,* and *B. holmesii* are respiratory pathogens that affect mammals, causing significant economic losses in the fields of human health and agriculture ^1^. Recent findings have identified *B. holmesii* as the second etiological factor of whooping cough. It does not produce most of the protein antigens of *B. pertussis*, such as PT, FHA, PRN, and FIM, and thus evades protection by the existing pertussis vaccines.

Bacterial pathogens expose on the surface a variety of complex carbohydrates that are essential for the structural integrity and interactions with hosts ^2–4^. Gram-negative bacteria produce lipopolysaccharides (LPS) which are the main components of the outer membrane of bacterial envelope. Besides they often produce several types of extracellular glycans vital for colonization and pathogenesis. Surface-bound capsular polysaccharides (CPS) ^5^ provide an extra protective layer at the surface, while some other bacteria can also synthesize exopolysaccharides (EPS), which are not anchored in the outer membrane, but instead are released into the environment. All these extracellular glycans play an important role in bacterial pathogenesis and survival. Capsular polysaccharides enable bacteria to evade the host immune responses as they provide shield against complement actions and hamper interactions with antibodies and macrophages. Therefore, capsules constitute important virulence factors.

The genomes of the *Bordetellae* contain loci that might encode a polysaccharide capsule. To date the references to *Bordetella* capsule-like glycans are scarce. However there is no clear indication what are the structural features of the capsules and these hypothetical capsules from the *Bordetellae* have neither been isolated nor structurally characterized so far ^6,7^. The genes involved in biosynthesis of the *Bordetella* capsules encode products that are homologous to the *Salmonella typhi* Vi antigen biosynthesis enzymes. Therefore the products of the *Bordetella* locus might resemble the Vi antigen, *i.e.* the polymer of the N-acetylgalactosaminuronic acid ^5,8^.

On the other hand, there are reports that *Bordetella* species produce an exopolysaccharide, namely the *Bordetella* polysaccharide (Bps). Immunological data suggest that Bps is a poly-b-1,6-D-Glc*p*NAc (PNAG) polysaccharide antigenically similar to this found in *Staphylococcus aureus* ^9^ and some Gram-negative bacteria ^10^. Bps is encoded by the bpsABCD operon and implicated in *Bordetella* biofilm formation ^11^. It is also claimed that Bps partakes in colonization of the respiratory tract and confers protection from complement-mediated bacterial killing ^12^. The capsule locus of *Bordetella* resembles those in other bacteria, however there is an inconsistency between *Bordetella* genome and the actual glycome encoded by this operon. To-date there is no clear confirmation of the structure of this capsular antigen and these glycan moieties produced by bacteria are elusive. It was suggested that *B. pertussis* may produce a microcapsule. Still no studies have ever reported on the isolation and structural features of the *B. pertussis* capsular glycan and whether the capsule is present among other *Bordetellae* ^13^. This makes the overall picture of the *Bordetella* glycome largely incomplete.

The efficacy of currently available pertussis vaccines is substantially lower than these of tetanus and diphtheria. Unlike tetanus and diphtheria infections for which all symptoms are caused exclusively by toxins, pertussis pathogenesis involves multiple virulence factors. Besides the secretory proteins of *B. pertussis* (e.g. pertussis toxin), surface molecules such as adhesins and endotoxin are involved in pathogenesis of pertussis. The bacterial extracellular polysaccharide of *B. bronchiseptica* has been implicated in facilitating transmission ^14^. Experimental studies using animals strongly indicate that current vaccines prevent symptoms and severe form of disease, but do not effectively prevent transmission of the bacterium from host to host. The exact role of the exo-glycans in not known.

Bps plays a pivotal role in maintaining the structural integrity of *Bordetella* biofilms under both *in vitro* and *in vivo* conditions ^11,12^. The formation of biofilms has been identified as a significant factor in the progression of *Bordetella pertussis* infections. However, the protective role of biofilm antigens and capsule-type glycans remains to be elucidated ^15^.

The pathogenesis of pertussis is complex, encompassing all stages from the initial infection through colonization and transmission ^16–19^. There is an absence of comprehensive data integration between genomic information and the resulting phenotypes concerning the full array of virulence factors implicated in the pathogenesis of whooping cough. Additionally, there are conflicting reports, regarding the existence and nature of the *Bordetella* exoglycans. Despite the existence of certain indications at the genomic level that these bacteria possess the capacity to produce capsules, no structural data regarding these molecules is currently available. In this paper, we describe the extracellular glycome elements of *Bordetellae* that were identified during our ongoing investigation into the elusive capsular antigen. We elucidate the structural features of the extracellular glycans and discuss their potential implications for the design of a novel pertussis vaccine.

## Results

### Isolation of the extracellular glycans of *Bordetellae*

The bacteria were removed from the media by centrifugation and supernatants were collected for the subsequent preparations of extracellular glycan-fractions. The supernatants were freeze-dried and subsequently fractionated using size-exclusion chromatography on a HiLoad Superdex 30 pg column. Three main fractions were obtained: fraction I (retention time range: 40-50), fraction II (retention time range: 51-69), and fraction III (retention time range: 70-80). The general pattern observed in the chromatograms for the exo-glycan fractions isolated from the culture media of different *Bordetella* species and strains were similar and differed only in the relative distribution of the components (**Fig. 1 A**). The fractions were analyzed by NMR spectroscopy, which indicated the glucose homopolymers in fraction I (**Supplementary Fig. 1**). Fraction II showed the same polysaccharide components, but with a lower degree of polymerization, whereas fraction III represented a complex oligosaccharide and fraction IV contained medium components and small metabolites. The oligosaccharide fraction III was the most abundant in all the analyzed *Bordetellae*, although the absolute amounts varied between species and strains (**Fig. 1 B**, **Supplementary Table 1**).

**Fig. 1.**
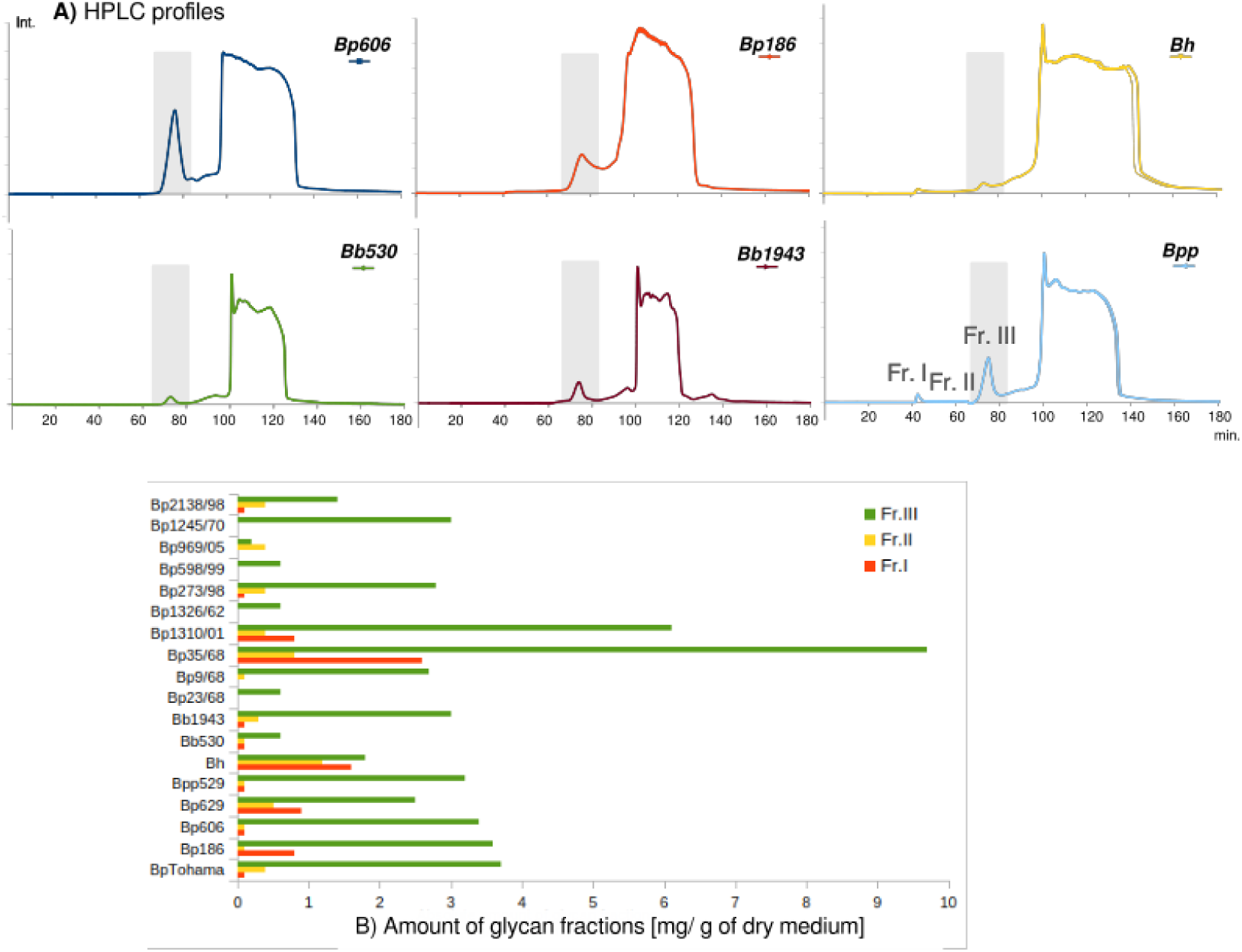
Isolation of exoglycans from the culture media of *Bordetellae*. A) Size-exclusion chromatography profiles of the exoglycans separated on the HiLoad Superdex 30 pg, with UV detection (190 nm). Profiles are shown for strains: *B. pertussis* 606 (*Bp*), *B. pertussis* 186, *B. holmesii* ATCC 51541 (*Bh*), *B. bronchiseptica* 530 (*Bb*), *B. bronchiseptica* 1943, and *B. parapertussis* 529 (*Bpp*). The indicated regions (gray boxes) represent the main poly- and oligosaccharide fractions (fr. III) used for the structural analysis. B) Distribution of the exoglycan fractions (Fr. I, Fr. II and Fr. III) recovered from cultures of the analyzed *Bordetella* species and strains [mg of glycans per 1g of dry post-culture medium].

### Structure analysis of the *Bordetellae* exooligosaccharides

The ^1^H (**Fig. 2**) and HSQC-DEPT (**Fig. 3 A, Supplementary Fig. 2**) NMR spectra, recorded for the main exooligosaccharide fraction (Fr. III), contained signals for up to nine anomeric protons and carbons. These signals derived from the main exooligosaccharide (*Bordetella* OligoSaccharide, BOS) as well as these from the short forms of glucose homopolymers.

**Fig. 2.**
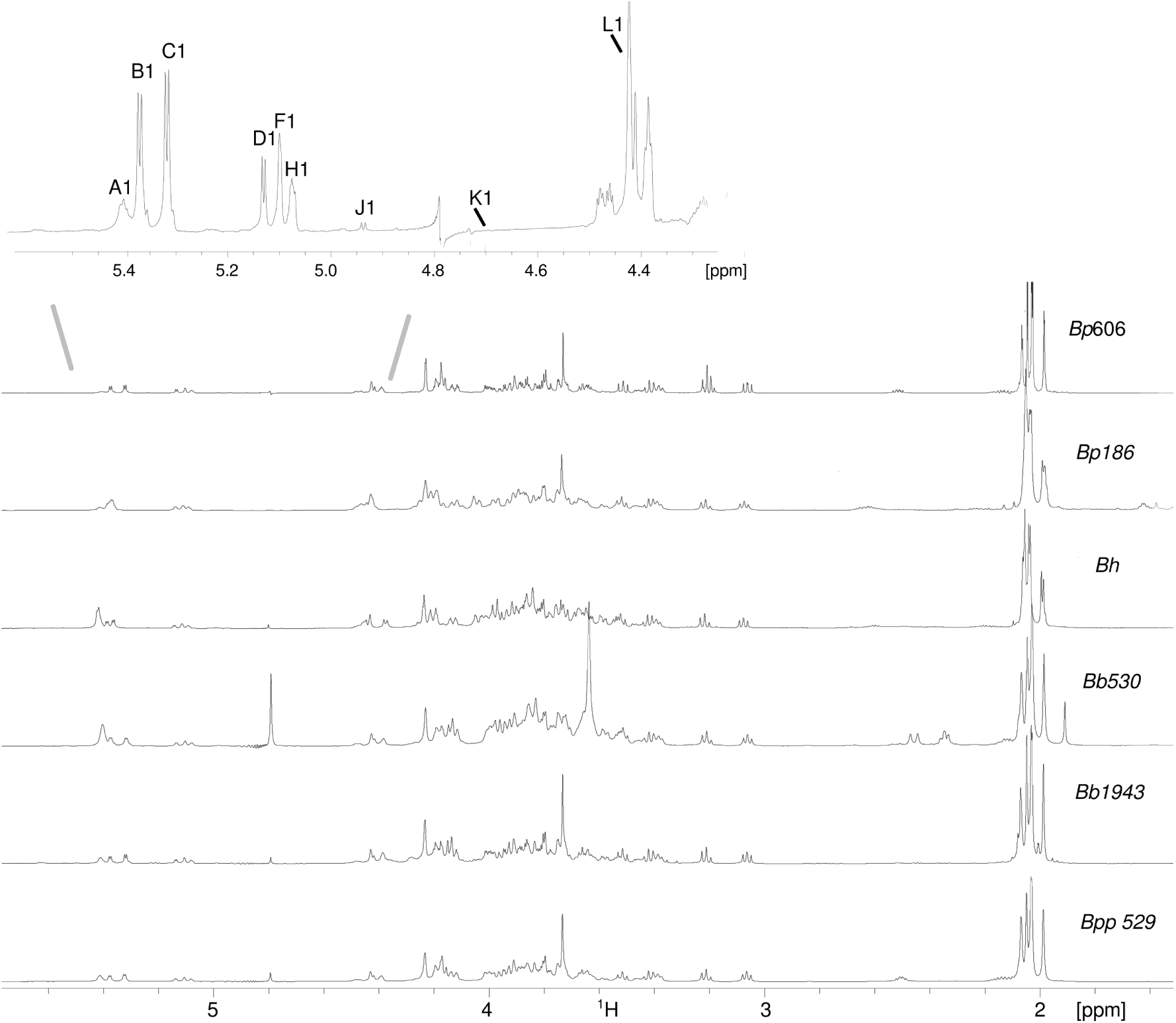
^1^H NMR spectra of the *Bordetella* oligosaccharides (fraction III). The region of anomeric proton signals is extended and the anomeric protons are assigned. The minor variants E1, G1, I1 are not assigned in the spectrum. The spectra were acquired for the OS isolated from the culture media of *B. pertussis* 606, *B. pertussis* 186, *B. holmesii ATCC 51541*, *B. bronchiseptica* 530, *B. bronchiseptica* 1943, and *B. parapertussis* 529. The spectrum was obtained for ^2^H_2_O solutions at 600 MHz and 25 °C.

**Fig. 3.**
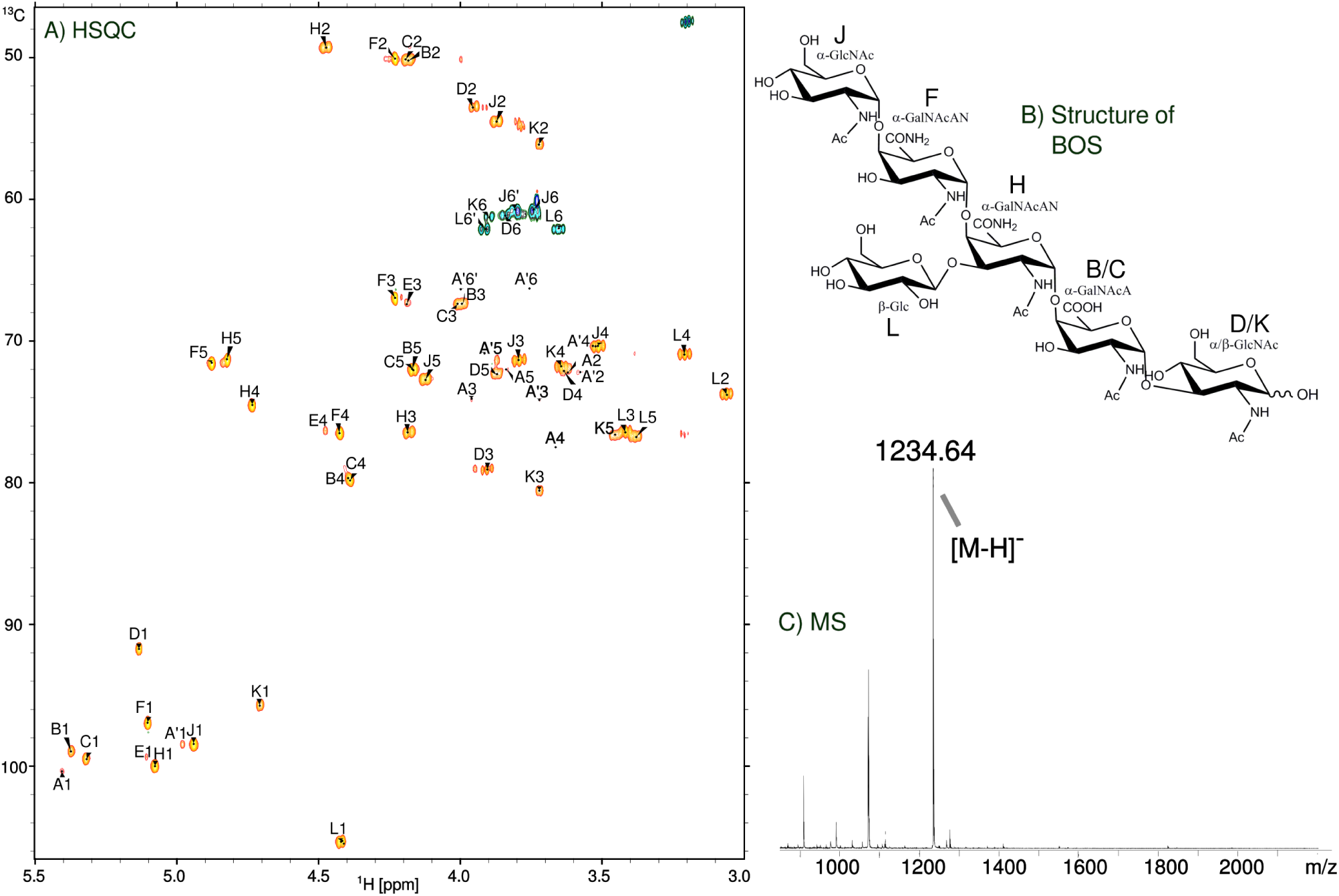
Structural analysis of *Bordetella* oligosaccharides (fraction III). A) HSQC-DEPT spectrum, in which the capital letters refer to carbohydrate residues as shown in the B) structure of the BOS. A and A’ residues correspond to the glucans. Residue D/K representing the →3)-α-Glc*p*NAc (**D**) and →3)-β-Glc*p*NAc (**K**) forms at the reducing end of a free hexasaccharide. C) The MALDI-TOF MS of the exooligosaccharide fraction recorded in the negative mode. The main ion corresponds to the BOS.

The major signals and spin-systems were assigned by COSY, TOCSY with different mixing times, HSQC-DEPT, HSQC-TOCSY and HMBC experiments. By comparing the chemical shifts with previously published NMR data for respective monosaccharides ^20,21^ and considering the ^3^J_H,H_-values for the coupling between ring protons, estimated from the cross-peaks in the two-dimensional spectra, the sugars were identified and their anomeric configuration determined. All the spin-systems comprising ^1^H and ^13^C resonances (**Table 1**) were determined.

**Table 1.** ^1^H and ^13^C NMR chemical shifts of the *Bordetellae* extracellular glycans ^a^.

| RESIDUE |  | Chemical shifts (ppm) |  |  |  |  |  |  |
| --- | --- | --- | --- | --- | --- | --- | --- | --- |
|  |  | H-1 | H-2 | H-3 | H-4 | H-5 | H-6a, 6b | H Ac |
|  |  | C-1 | C-2 | C-3 | C-4 | C-5 | C-6, CO | C Ac (CO, CH<sub>3</sub>) |
| **A** | → 4)-α-Glcp-(1 → • | 5.41 | 3.64 | 3.98 | 3.67 | 3.85 | 3.83, 3.86 |  |
|  |  | 100.4 | 72.4 | 74.2 | 77.5 | 71.9 | 61.2 |  |
| **A'** | → 6)-α-Glcp-(1 → • | 4.98 | 3.59 | 3.72 | 3.53 | 3.91 | 3.75, 3.99 |  |
|  |  | 98.5 | 72.4 | 74.0 | 70.3 | 71.0 | 66.3 |  |
| **B** | → 4)-α-GalpNAcA-(1 → | 5.38 | 4.20 | 4.02 | 4.42 | 4.28 | - | 2.03 |
|  |  | 0.004 | 0.008 | 0.016 | 0.026 | 0.101 |  | 0.002 |
|  |  | 98,9* | 50.2 | 67.4 | 79.6 | 71.9 | 174.3 | 175.5, 22.8 |
|  |  | 0,32 | 0.31 | 0.43 | 0.56 | 0.70 | 0.3 | 0.32 |
| **C** | → 4)-α-GalpNAcA-(1 → | 5.34 | 4.18 | 4.00 | 4.41 | 4.28 | - | 2.03 |
|  |  | 0.018 | 0.007 | 0.013 | 0.027 | 0.101 |  | 0.002 |
|  |  | 99.5 | 50.2 | 67.4 | 79.8 | 72.0 | 174.3 | 175.5, 22.8 |
|  |  | 0.31 | 0.29 | 0.43 | 0.53 | 0.63 | 0.3 | 0.32 |
| **D** | → 3)-α-GlcpNAc | 5.13 | 3.96 | 3.90 | 3.63 | 3.86 | 3.82 | 1.98 |
|  |  | 0.002 | 0.006 | 0.010 | 0.010 | 0.011 | 0.006 | 0.002 |
|  |  | 91.7 | 53.5 | 79.1 | 72.1 | 72.3 | 61.1 | 175.1, 22.8 |
|  |  | 0.32 | 0.29 | 0.41 | 0.36 | 0.19 | 0.27 | 0.32 |
| **E^** | → 4)-α-GalpNAcAN-(1 → | 5.10 | 4.02 | 4.18 | 4.48 | - | - | - |
|  |  | 0.010 | 0.015 | 0.012 | 0.002 |  |  |  |
|  |  | 99.4 | 50.1 | 67.4 | 76.3 | - | - | - |
|  |  | 0.36 | 0.05 | 0.36 | 0.36 |  |  |  |
| **F^** | → 4)-α-GalpNAcAN-(1 → | 5.10 | 4.23 | 4.23 | 4.43 | 4.87 | - | 2.03 |
|  |  | 0.002 | 0.002 | 0.002 | 0.002 | 0.009 |  | 0.002 |
|  |  | 96.9 | 50.1 | 66.9 | 76.5 | 71.5 | 174.4 | 174.9, 22.9 |
|  |  | 0.32 | 0.31 | 0.31 | 0.33 | 0.33 | 0.3 | 0.32 |
| **H** | → 3,4)-α-GalpNAcAN-(1 → | 5.08 | 4.46 | 4.20 | 4.74 | 4.83 | - | 2.07 |
|  |  | 0.002 | 0.009 | 0.006 | 0.004 | 0.005 |  | 0.002 |
|  |  | 100.0 | 49.3 | 76.5 | 74.5 | 71.3 | 174.6 | 175.7, 23.2 |
|  |  | 0.34 | 0.30 | 0.36 | 0.35 | 0.35 | 0.3 | 0.32 |
| **J** | α-Glc<sub>p</sub>NAc-(1 → | 4.94<br>0.002 | 3.88<br>0.008 | 3.79<br>0.004 | 3.51<br>0.005 | 4.12<br>0.003 | 3.73, 3.75<br>0.003, 0.007 | 2.04<br>0.002 |
|  |  | 98.4<br>0.32 | 54.5<br>0.32 | 71.3<br>0.33 | 70.3<br>0.33 | 72.7<br>0.30 | 60.7<br>0.33 | 174.8, 22.9<br>0.32 |
| **K** | → 3)-β-Glc<sub>p</sub>NAc | 4.71<br>0.005 | 3.72<br>0.005 | 3.71<br>0.006 | 3.66<br>0.003 | 3.46<br>0.002 | 3.85<br>0.009 | 1.98<br>0.002 |
|  |  | 95.7<br>0.33 | 56.1<br>0.26 | 80.6<br>0.44 | 71.8<br>0.33 | 76.6<br>0.35 | 61.3<br>0.30 | 175.5, 22.8<br>0.32 |
| **L** | β-D-Glc<sub>p</sub>-(1 → | 4.43<br>0.003 | 3.06<br>0.004 | 3.42<br>0.002 | 3.21<br>0.002 | 3.39<br>0.001 | 3.66, 3.91<br>0.007, 0.009 | - |
|  |  | 105.3<br>0.33 | 73.8<br>0.32 | 76.5<br>0.31 | 71.0<br>0.32 | 76.8<br>0.31 | 62.0<br>0.35 | - |

<sup>a</sup>The chemical shifts are given as averaged values for the residues in the same environment obtained from the spectra acquired for the exooligosaccharide fractions of *B. holmesii* ATCC 51541, *B. pertussis* strains 186 & 606, *B. parapertussis* 529, *B. bronchiseptica* strains 530 and 1943 except from the residues A and A' as these were determined for the most abundant fraction I of *B. holmesii* (**Supplementary Fig. 2**).
The values marked with (^) represent the same residue → 4)-α-GalpNAcAN-(1 → in a different microenvironment. All assignments were performed with a NMRFAM-SPARKY software. The standard deviation values are printed in a small font-type. All spectra were obtained for <sup>2</sup>H<sub>2</sub>O solutions at 25 °C. Acetone (δ<sub>H</sub> 2.225, δ<sub>C</sub> 31.05) was used as internal reference.
The residues constituting the fraction I (glucan) are marked with (•). For complete assignment see **Supplementary Fig. 1**.
The absolute configuration was determined as D-glucopyranose for the residues D, J, K and L, using the method of York *et al.* <sup>22</sup>.

Residue **A** with the H-1/C-1 signals at δ 5.41/100.4, *J*_H-1,H-2_ 3.4 Hz, was assigned as the 4-substituted a-Glc*p* on basis of the large vicinal coupling constants between H-2, H-3, H-4 and H-5 protons, and the downfield shift of the C-4 signal (δ 77.5 ppm). Similarly, residue **A’** with the H-1/C-1 signals at δ 4.98/98.5 ppm, J_H-1,H-2_ 3.3 Hz was assigned as the 6-substituted a-D-Glc*p* on basis of the downfield shift of the C-6 signal (δ 66.3 ppm). The **A** and **A’** residues constitute the a-(1→4)-linked and a-(1→6)-linked homopolymers of glucose that in the long-chain forms were identified previously in the fraction I (**Supplementary Fig. 1**). The fraction III is a mixture of these glucans with the more complex oligosaccharide comprising the following residues **B** – **L**.

Residue **B** with the H-1/C-1 signals at δ 5.38/98.9 ppm, *J*_H-1,H-2_ 3.9 Hz, was assigned as the 4-substituted a-Gal*p*NAcA residue based on the characteristic five proton spin system, the low chemical shift of the C-2 signal (δ 49.3) the high chemical shifts of H-4 (δ 4.42), H-5 (δ 4.28), and the C-6 (δ 174.3) signals and the small vicinal couplings between H-3, H-4 and H-5.

Residue **C** with the H-1/C-1 signals at δ 5.34/99.5 ppm, *J*_H-1,H-2_ 3.9Hz, was assigned as the 4-substituted a-Gal*p*NAcA residue based on the characteristic five proton spin system, the high chemical shifts of H-4 (δ 4.41), H-5 (δ 4.28), and the C-6 (δ 174.3) signals. The C residue is a variant of the residue B.

Residue **D** with the H-1/C-1 signals at δ 5.13/91.7 ppm, *J*_H-1,H-2_ 3.5 Hz, was assigned as the 3-substituted a-D-Glc*p*NAc residue from the low chemical shifts of the C-2 signal (δ 53.5 ppm), the relatively high chemical shift of C-3 (δ 79.1 ppm), and the strong vicinal couplings between H-2, H-3, H-4 and H-5.

Residue **F** with the H-1/C-1 signals at δ 5.10/96.9 ppm, *J*_H-1,H-2_ 2.5 Hz, was assigned as the 4-substituted a-Gal*p*NAcAN residue based on the characteristic five proton spin system, the low chemical shift of the C-2 signal (δ 50.1 ppm), the relatively high chemical shift of C-4 (δ 75.6 ppm) the high chemical shifts of H-4 (δ 4.43), H-5 (δ 4.87), and the C-6 (δ 174.4) signals and the small vicinal couplings between H-3, H-4 and H-5. Residue **E** with the H-1/C-1 signals at δ 5.10/99.4 ppm represents a minor variant of residue **F**.

Residue **H** with the H-1/C-1 signals at δ 5.08/100.0 ppm, *J*_H-1,H-2_ < 2 Hz, was assigned as the 3,4-disubstituted a-Gal*p*NAcAN residue based on the characteristic five proton spin system, the low chemical shift of the C-2 signal (δ 49.3 ppm), the relatively high chemical shifts of C-3 (δ 76.5 ppm) and C-4 (δ 74.5 ppm) signals, the high chemical shifts of H-4 (δ 4.74), H-5 (δ 4.83), and the C-6 (δ 174.6) signals and the small vicinal couplings between H-3, H-4 and H-5.

Both residues **F** and **H** were amidated at the carbonyl group as indicated by the combination of data from 1D ^1^H NMR run in H_2_O/D_2_O, the complementary ^1^H,^15^N-HSQC (identification of individual amide protons of -NH_2_, **Supplementary Fig. 3**) and ^1^H,^13^C-HMBC spectrum (linkage between one of the -NH_2_ protons to carbon C-5). This observation was further supported by a lack of pH-dependence of the H-5 chemical shifts in the HSQC-DEPT spectra acquired at pH ∼5 and pH ∼9.

Residue **J** with the H-1/C-1 signals at δ 4.94/98.4 ppm, *J*_H-1,H-2_ 3.6 Hz, was assigned as the terminal a-D-Glc*p*NAc residue based on the low chemical shifts of the C-2 signal (δ 54.5 ppm) and the strong vicinal couplings between H-2, H-3, H-4 an H-5.

Residue **K** with the H-1/C-1 signals at δ 4.71/95.7 ppm, *J*_H-1,H-2_ 7.0 Hz, was assigned as the 3-substituted b-D-Glc*p*NAc residue from the low chemical shifts of the C-2 signal (δ 56.1 ppm), the relatively high chemical shift of C-3 (δ 80.6 ppm), and the large vicinal couplings between all ring protons.

Residue **L** with the H-1/C-1 signals at δ 4.43/105.3 ppm, *J*_H-1,H-2_ 8 Hz, was recognized as the terminal b-D-Glc*p* from the similarity of the ^1^H and ^13^C chemical shifts and the large vicinal couplings between all protons as in the monosaccharide b-D-Glc*p*.

The absolute configurations of the sugars were determined according to method described by York *et al.* ^22^ using ^1^H NMR analysis of the *per-O*-(S)-2-methylbutyrate derivatives. Determination of the absolute configuration revealed the presence of D-*gluco*-pyranose configuration for the residues D, J, K and L.

The ^1^*J*_C-1,H-1_ values, obtained from an HSQC experiment run without carbon decoupling, confirmed the a-pyranosyl configuration for residues **B/C** (177 Hz)**, D** (173 Hz), **F** (178 Hz), **H** (178 Hz) and **J** (178 Hz), and b-pyranosyl configuration for residues **K** (165 Hz) and **L** (164 Hz).

The inter-residue connectivities between adjacent sugar residues were acquired by NOESY and HMBC experiments (**Supplementary Fig. 4, Table 2**). The results identified all disaccharide elements in the exooligosaccharides and thus provided the sequence of monosaccharides in the oligosaccharide (**Fig. 3**). For BOS inter-residue NOEs were found between H-1 of **J** and H-4 of **F**, H-1 of **F** and H-4 of **H**, H-1 of **L** and H-3 of **H,** H-1 of **H** and H-4 of **B**, H-1 of **B** and H-3 of **D/K**. Residue D/K constituted a free-reducing end of the oligosaccharide and gave rise to the **B** and **C** variants of this disaccharide segment. The HMBC spectra showed cross-peaks between the anomeric proton and the carbon at the linkage position and between the anomeric carbon and the proton at the linkage position (**Table 2**), which confirmed the sequence of sugar residues in the hexasaccharide. Thus the combined results indicate the following structure of the new exooligosaccharide (**Fig. 3 B**).

**Table 2.** Selected inter-residue NOE and ^3^*J*_H,C_-connectivities from the anomeric atoms of the exooligosaccharide of *Bordetellae*.

| Residue | Atom<br>$\delta_H/\delta_C$ (ppm) | Connectivities to | | Inter-residue<br>atom/residue |
| --- | --- | --- | --- | --- |
| | | $\delta_C$ | $\delta_H$ | |
| **B** → 4)-α-GalpNAcA-(1 → | H-1/C-1<br>5.38/98.9 | 80.6 | 3.71 | **C-3, H-3 of K** |
| **C** → 4)-α-GalpNAcA-(1 → | 5.34/99.5 | 79.1 | 3.90 | **C-3, H-3 of D** |
| **D** → 3)-α-Glc pNAc | 5.13/91.7 | - | - | **-\*** |
| **E** → 4)-α-GalpNAcAN-(1 → | | | | |
| **F** → 4)-α-GalpNAcAN-(1 → | 5.10/96.9 | 74.5 | 4.74 | **C-4, H-4 of H** |
| **H** → 3,4)-α-GalpNAcAN-(1 → | 5.08/100.0 | 79.6 | 4.42 | **C-4, H-4 of B** |
|  |  | 79.8 | 4.41 | **C-4, H-4 of C** |
| **J** α-Glc pNAc-(1 → | 4.94/98.4 | 76.5 | 4.43 | **C-4, H-4 of F** |
| **K** → 3)-β-Glc pNAc | 4.71/95.7 | - | - | **-\*** |
| **L** β-D-Glc p-(1 → | 4.43/105.3 | 76.5 | 4.20 | **C-3, H-3 of H** |

<sup>a</sup>The chemical shifts are given as averaged values for the residues in the same environment. The values marked with (\*) represent $\alpha$ and $\beta$ forms of the residue constituting the oligosaccharide reducing end.

On the basis of the structural data derived from the NMR experiments the theoretical monoisotopic mass of the BOS was calculated as m/z 1234.43 Da. The MALDI-TOF mass spectra of the exooligosaccharide fraction (BOS) recorded in the negative mode showed the main ion of monoisotopic molecules [M-H]^-^ at m/z 1234.64 (**Fig. 3 C**). The ESI MS spectrum confirmed the molecular size of the BOS (**Supplementary Fig. 5**). This ion corresponds to a BOS hexasaccharide. Thus the combined results indicate the following structure of the new exooligosaccharide (**Fig. 3 B**).

### Metabolic labeling of the *Bordetella* oligosaccharide

To confirm the biosynthetic origin of the BOS exoglycan we used *B. pertussis* isolate strain 35/68 for a small scale culture with a primary carbon source - sodium L-glutamate in the chemically defined Steiner-Scholte medium replaced by the ^13^C,^15^N-L-glutamate. The size-exclusion chromatography profile of the post-culture supernatant sample on the HiLoad Superdex 30 pg column revealed the pattern identical to this observed in the chromatograms for the *Bordetella* exo-glycan. The fractions were analyzed by NMR spectroscopy. 1D ^13^C and HSQC spectra of the metabolically labeled BOS confirmed that it was biosynthetically produced by *B. pertussis* strain (**Fig. 4**).

**Fig. 4.**
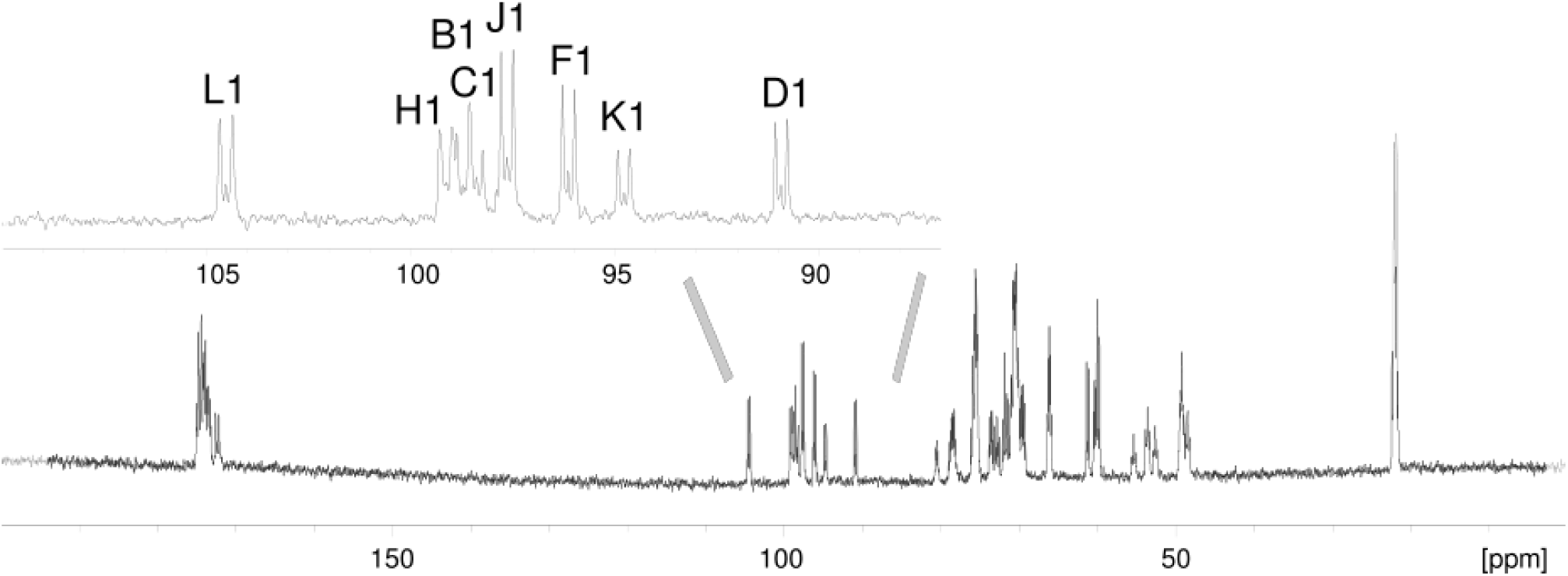
1D ^13^C NMR spectrum of ^13^C,^15^N-enriched *Bordetella* oligosaccharide BOS. The region of anomeric carbons is extended. The capital letters refer to carbohydrate residues and the numbers refer to carbons in the respective residue as shown on the structure in the Fig. 3.

### Reactivities of anti-*B. pertussis* antibodies with BOS

The BOS was identified as a new abundant exocellular component of the *Bordetellae* glycome. However, the antigenic properties and possible roles of this glycan in the interactions with hosts remain unknown. As no antibodies specific to this glycan were available, we used anti-pertussis sera from our collection (**Supplementary Table 2**) to test for the possible cross-reactivities. In the study we included polyclonal antibodies against (1) the whole bacterial cells (WB) of various strains of *B. pertussis* (anti–*Bp* 186, anti–*Bp* mix of strains 186, 576, 606 and 629), and (2) the neoglycoconjugates of *B. pertussis* lipooligosaccharide (LOS)-derived glycans (anti–disaccharide-HSA, anti–OS-PT, anti–pentasaccharide-PT, anti–pentasaccharide-TTd). Additionally, the reactivities of antibody against antigen Vi (anti-Vi, commercial Ab) were tested. Antibodies from the unrelated serum were used as control. The cross-reactivities of these antibodies with BOS were determined by immunoblotting (**Fig. 5 A**). All antibodies against the whole bacterial cells reacted with BOS in the dot-blot screening. Similar reactions were also observed for the anti-conjugate antibodies although the intensities differed. The reactions could be explained by the presence of the cross-reacting epitopes comprising the terminal Glc*p*NAc residue at the non-reducing in the LOS of the *B. pertussis* bacterial cells and the LOS-derived distal glycans – OS, disaccharide and pentasaccharide within the conjugates. Only weak binding of the BOS with the anti-Vi antibodies was detected, confirming that these two glycans are immunologically diverse and belong to different types of molecules present in *Bordetellae*.

**Fig. 5.**
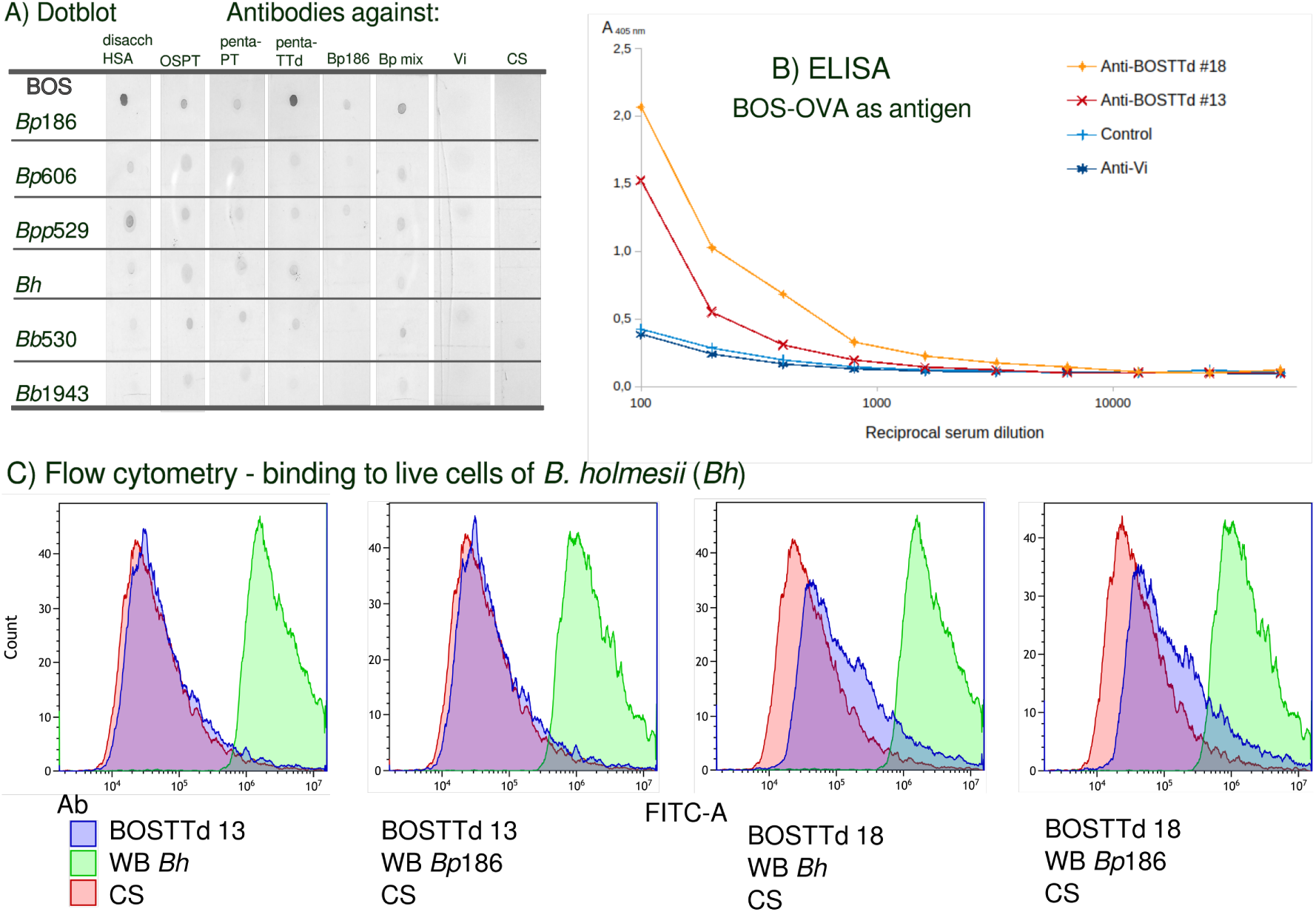
Serological characterization of *Bordetella* exooligosaccharide BOS. A) Reactivities of the isolated exoglycans with polyclonal antibodies against *B. pertussis* whole bacterial cells (*Bp*186 and *Bp* mix) and LOS-derived antigens determined by dot-blotting. Each row represents exoglycans BOS isolated from the respective *Bordetella* species/strains. The antibodies used are indicated in the header. The anti-conjugate antibodies were raised against LOS-derived dodecasaccharide (*Bp*186 OS) and its isolated segments containing distal epitope at the non-reducing residue (disaccharide – disacch, pentasaccharide – penta). HSA – human serum albumin, PT – pertussis toxin, TTd – tetanus toxoid, Vi – Vi antigen, CS – control serum; B) Reactivities of polyclonal antibodies against BOS-TTd conjugate with BOS-OVA conjugates used as solid-phase antigens in ELISA. Antibodies from two different rabbits (#13 and #18) were tested. Anti-Vi antibodies and a pre-immune serum were used as control. The ELISA values are the means of three replicates. The standard deviations did not exceed 5% and are not shown. C) Binding of anti-BOS-TTd conjugate and anti-whole-bacteria antibodies to live cells of *B. holmesii* strain ACTC 51541 evaluated by flow cytometry. The histograms show the results for control serum (CS), anti-BOS-TTd conjugate sera, and sera against whole cells of *B. holmesii* and *B. pertussis* 186 (WB).

### BOS neoglycoconjugates and reactivities of anti-BOS-TTd antibodies

Immunochemically, a free BOS is a hapten, therefore to obtain the BOS specific antibodies we converted it to an immunogenic form by covalent coupling to a protein carrier to form a neoglycoconjugate. We followed the standard procedure of selective oxidation at the reducing end of the oligosaccharide (residue D/K). A single aldehyde group formed in the BOS (**Supplementary Fig. 6**) was used to establish the covalent linkage with a monomeric fraction of tetanus toxoid by reductive amination ^23,24^. The BOS-TTd neoglycoconjugate was used for immunization of rabbits. The second batch of BOS was used to synthesize the complementary conjugate with the OVA protein carrier. The BOS-OVA conjugate was used as a solid state antigen in the detection system for the anti-BOS-TTd antibodies. The reactivity of anti-BOS-TTd conjugate antibodies with BOS-OVA was tested in ELISA assay. The end point titers in ELISA (**Fig. 5 B**) and a distinct reaction of anti-conjugate antibodies with the BOS-OVA in dot-blotting showed that the conjugate was immunogenic in rabbits and that the antibodies were capable of recognizing the BOS.

To investigate the presence of the BOS on the bacterial cells in addition to the exoglycan form the binding of anti-BOS-TTd antibodies with whole bacterial cells was tested, using flow cytometry. *B. holmesii* strain ATCC 51541 bacteria were incubated with anti-BOS-TTd antibodies and antibodies against the whole bacterial cells (anti-*B. holmesii*, anti-*B. pertussis* 186). Pre-immune serum was used as a negative control. FACS analysis showed that anti-WB *B. holmesii* and anti-WB *B. pertussis* 186 antibodies reacted strongly and in a similar pattern with live cells of *B. holmesii* (**Fig. 5 C**). Whereas the histograms (FL1-H) for the reaction of the anti-BOS-TTd antibodies with bacterial cells showed that the fluorescence intensity was similar to the control.

Only a small population of bacteria was labeled with the higher intensity when the Ab#18 was used. This could be explained by the presence of a minor population of BOS molecules associated with the outer membrane or alternatively by the cross-reacting epitope recognized in the LPS, comprising the terminal Glc*p*NAc residue at the non-reducing end. FACS analysis indicated that the anti-BOS-TTd antibodies labeled bacteria with lower intensity than anti-WB antibodies and on par with the control pre-immune serum. This observation confirms that the prevailing form of BOS is the exoglycan released to medium during the planctonic growth of *Bordetellae*.

## Discussion

In our investigations of a complete *Bordetella* glycome, we discovered the new genus specific extracellular oligosaccharide. As it was identified among all the investigated *Bordetella* species and strains we named it the *Bordetella* oligosaccharide (BOS). Unlike other extracellular glycans, which are typically polysaccharides (capsules, *Bps*, EPS, ECA) BOS is a free hexassaccharide. It was first recovered from the chemically defined Steiner-Scholte medium which is routinely used to grow *B. pertussis* in planctonic cultures.

Pertussis pathogenesis involves multiple virulence factors such as secretory proteins, surface adhesins and lipooligosaccharides (LOS). To date references that indicate the presence of glycans among *Bordetella* species are scarce and mostly limited to these that are the integral part of the cell outer membrane (LOS, LPS) as well as these attached to the cell (Bps). The possible encapsulation of *Bordetella* species was partly deduced from the polysaccharide staining techniques. The capsule detection procedures have led to a conclusion that *Bordetella pertussis* may produce a *micro-capsule* ^13^. However, the structures of the these glycans encoded and produced by *Bordetellae* remain elusive.

The hypothetical capsules and exopolysaccharides of *Bordetellae* have not been isolated and structurally characterized so far ^7^. Moreover, reports on structures of these glycans are inconsistent. Some authors claim that genes involved in biosynthesis of *Bordetella* capsules encode products homologous to *Salmonella typhoid* Vi antigen ^8^. Contrary reports suggest that *Bordetella spp.* produce PNAG-like exopolysaccharide (*Bps*) ^9^, similar to this of *Staphylococcus aureus* ^10^ and a conserved structure produced by major bacterial, fungal and protozoal parasites ^25^. Bps was implicated in biofilm formation and colonization of the respiratory tract. It confers protection from complement-mediated bacterial killing ^12,26,27^. These exopolysaccharides are required for *in vitro* biofilm development ^2^ and are involved in the formation/stabilization of the three-dimensional architecture of mature biofilms ^11^. There are claims that the Bps is antigenically similar to the PNAG-type poly-b-1,6-D-Glc*p*NAc polysaccharides produced by diverse bacterial species ^10,28^. It has been implicated in colonization of the respiratory tract and protection from complement-mediated bacterial killing ^12,26,27^. And yet, no attempts at determining the chemical structures of the Bps were actually reported. More importantly, the previous studies indicated, that Bps was predominantly anchored to the bacterial cells, with only a small fraction released during planctonic cultures of *B. pertussis* ^29^. Our attempts to isolate the released form of Bps from the post-culture media of different *Bordetella* species failed. Instead we have identified a new type of extracellular glycans - *Bordetella* oligosaccharide. BOS was predominantly released in the culture media. FACS analysis indicated no interactions of the anti-BOS-TTd antibodies with the whole bacteria and thus supported the observation that BOS was an exo-oligosaccharide.

The biosynthetic origin of the BOS was confirmed by nuclear magnetic resonance (NMR) combined with metabolic labeling in culture, using ^13^C,^15^N-L-glutamate as a primary carbon source. The labeled BOS revealed the chemical shifts of all the residues virtually identical to that for the BOS isolated from a standard culture.

The role this molecule may play in the interactions with the host is unclear. As the *Bordetella* oligosaccharide is a relatively small molecule, it constitutes a hapten. To date nothing is known about the forms BOS is presented to the host immune system by bacteria, or whether these forms are immunogenic. In this studies we showed that anti-*B. pertussis* antibodies, specific towards defined surface related glycans of *B. pertussis* reacted strongly with the isolated BOS. However, the cross-reactivities of the set of *B. pertussis* anti-neoglycoconjugate antibodies could be explained by the presence of common structural motifs among the LOS and LPS of *Bordetellae*. We demonstrated previously ^30^ that the terminal GlcNAc is an immunodominant component of *B. pertussis* 186 LOS. The GlcNAc residue is also a constituent of the BOS, therefore it might be recognized by monospecific anti-disacch-HSA, anti-penta-PT, anti-penta-TTd, anti-OS-PT, and anti-OS-TTd antibodies. Previously it was suggested ^13^, that *B. pertussis* produces a micro-capsular polysaccharide composed of N-acetylgalacturonic acid, but such Vi-like micro-capsule was never isolated. We demonstrated that anti-Vi antibodies showed only weak cross-reactions with BOS. This could be explained by the presence of Gal*p*NAcA residues in the BOS structure, conformationally similar to N-acetylgalacturonic acid component of Vi antigen.

A potential consequence of BOS release is the disruption of the interactions between bacteria and the host immune system, which could have significant implications for immunological processes. BOS could prevent anti-LPS antibodies from reaching the target by introducing an interference with the opsonization of bacteria. Consequently, BOS has the potential to circumvent the host’s defense mechanisms. Similarly, the presence of the BOS in the preparations of glycans could affect the interpretation of the observed immunological properties of Bps. We believe that understanding of *Bordetella* glycome could resolve the problem of missing elements in the composition of the pertussis vaccine that contribute to its largely incomplete efficiency.

## Methods

### *Bordetella* strains culture

The bacterial strains of genus *Bordetella* were obtained from: the National Medicines Institute (Warsaw, Poland), the Polish Collection of Microorganisms (Hirszfeld Institute of Immunology and Experimental Therapy, Polish Academy of Sciences, Wrocław, Poland), the German Collection of Microorganisms and Cell Cultures (Leibniz-Institut, Berlin, Germany) and from National Collection of Type Cultures (UK Health Security Agency Salisbury, United Kingdom). All strains are listed in Supplementary Table S3 (**Supplementary Table S3**). Bacteria were reconstituted from the short-term collection (local stock, *i.e.* bacterial suspension in the Steiner-Scholte medium, containing 20% glycerol, stored at -75 °C) and cultured at 37 °C for 3 days on a charcoal agar (Oxoid, United Kingdom) plates containing 10% the defibrinated sheep blood (GRASO Biotech, Owidz, Poland). Bacteria collected from the plates were used to prepare liquid inocula (∼250 mL) in the Steiner-Scholte medium containing 10% of growth factors for the liquid culture (3-5 days at 37 °C) at the larger scale (composition of the media in **Supplementary Table 4**). For the metabolically ^13^C,^15^N-labeled small scale production of exoglycans the bacteria were cultured in the modified Steiner-Scholte medium. In the modified medium the main carbon source -L-glutamate was replaced by the ^13^C,^15^N-labeled L-glutamate. The cultures were aerated by shaking (Infors HT Ecotron, 140 rpm, 3-5 days at 37°C). The bacteria were harvested from the medium by centrifugation 8000 × g, 30’, 4°C (Sorvall Lynx 6000) and supernatants were collected for the subsequent preparations of EPS-fractions. The purity of bacterial cultures was monitored by biotyping using MALDI-TOF MS Biotyper method (MBT).

### Isolation of *Bordetella* extracellular glycans

The lyophilized supernatants obtained from the bacterial culture were fractionated using the semi-preparative HPLC UltiMate 3000 chromatographic system (Dionex Corporation, Sunnyvale, CA, USA) on HiLoad 16/600 Superdex 30 prep grade column (30 mm × 124 cm, grain size 34 μm, GE Healthcare, Chicago, IL, USA) equilibrated with 0.05 M acetic acid. The eluates were monitored spectrophotometricaly and by differential refractometry using an UV detector at wavelengths λ = 190 nm, λ = 206 nm and λ = 280 nm and a Shodex RI-102 detector (Showa-Denko Tokyo, Japan). All fractions were checked by NMR spectroscopy and MALDI-TOF mass spectrometry. Polysaccharides, oligosaccharides and their reduced and oxidized forms were purified on a TSKgel G3000PW column (21.5 mm × 60 cm, grain size 12 µm, Tosoh Bioscience, Japan) equilibrated with mQ water. A TSKgel G-3000SW column (21.5 mm × 30 cm, grain size 13 μm, Tosoh Bioscience, Japan) equilibrated with PBS was used to purify glycoconjugates and tetanus toxoid.

### GC-MS analyses

Monosaccharides were analyzed as their alditol acetates by GC-MS using a Thermo ITQ™ 1100 GC-Ion Trap mass spectrometer coupled with a Trace™ 1310 gas chromatograph (Thermo Scientific™) equipped with HP-5M S column (30 m, ID = 0.25 mm, dF = 0.25 μm) (Agilent Technologies, Lexington, MA, USA) and a temperature gradient 150–270 °C at 8 °C·min^−1^. Helium was used as the carrier gas at a constant flow of 1.8 ml/min (temperature program 150°C–270°C, 8 °C/min). The absolute configurations of the sugars were determined using ^1^H NMR spectroscopy by converting the polysaccharide hydrolysate components and relevant monosaccharide standards into O-(S)-2-methyl butyrate derivatives as described by York *et al.* ^22^. For NMR analyzes, the samples were dissolved in acetone-d5 (∼160 μL) and transferred to 3 mm diameter NMR tubes.

### MS analyses

MALDI-TOF mass spectrometry analyses were carried out using the Ultraflex instruments (Bruker Daltonics, Germany). Spectra were acquired in linear and reflectron modes, with negative and positive ion polarization. 2,5-dihydroxybenzoic acid (DHB) was employed as matrix. Spectra were calibrated over the mass range from ∼1000 to 3500 m/z using the Peptide Calibration Standard (Bruker Daltonics). Electrospray ionization (ESI) MS measurements were performed on an amaZon SL device (Bruker Daltonics, Germany) equipped with an ion trap detector. The analyzed oligosaccharides were dissolved in an acetonitrile/water mixture (1:1). The sample concentration was 50 µg/ml. Spectra were recorded in the range m/z 200-2000 (measurements in negative ion mode). The instrument was calibrated using ESI-L Tuning Mix reagent (Agilent Technologies, USA) in positive ion mode. The following parameters were used for the ion source: sample flow 3 µl/min, source temperature 200°C, carrier gas flow (nitrogen) 5 L/min at a pressure of 8 psi.

### NMR spectroscopy

All NMR spectra were obtained for ^2^H_2_O and H_2_O/ ^2^H_2_O [90%/10%] solutions with a Bruker Avance III 600 MHz spectrometer (Bruker). Spectra were recorded using QCI Cryoprobe, at 25 °C and the excitation sculpting pulse sequence was used for the suppression of the residual HOD signal. The basic data sets were provided by observation of carbohydrate-relevant nuclei, ^1^H, ^13^C, ^15^N and ^31^P in various combinations, comprising COSY, TOCSY, NOESY, HSQC-DEPT, HSQC-TOCSY and HMBC experiments. The data were acquired and processed using standard Bruker software (Topspin). The processed spectra were assigned using the NMRFAM-SPARKY program ^31^.

### Preparation of neoglycoconjugates

*(1) BOS-TTd conjugate.* The isolated BOS was oxidized with periodate solution (NaIO_4_). The oxidized BOS was subsequently purified by HPLC using molecular sieve chromatography on the TSKgel G3000-PW column (2.1 cm × 30 cm) equilibrated with water. The presence of an active aldehyde group in the oxidized compound was verified by NMR spectroscopy. The selectively oxidized fraction, bearing an active aldehyde group was conjugated to a tetanus toxoid by the reductive amination as described previously ^23,24^. Glycoconjugate was fractionated on a G3000-SW column equilibrated with phosphate-buffered saline (PBS, pH 7.5). Fractions containing the conjugate were concentrated by ultrafiltration and stored at 4 °C with an addition of thiomersal (0.01%).
*(2) BOS-OVA conjugate.* The isolated and purified BOS was first conjugated to an adipate dihydrazide (ADH). The conjugation was carried out as described previously ^32^. Briefly, the oligosaccharide (15 mg) solution in H_2_O (100 μl) was mixed with a 10% solution of ADH in H_2_O, and the mixture stirred at ∼22 °C for 72h. It was than followed by neutralization with NaHCO_3_ and purified by HPLC (TSKgel G3000-PW 2.1 cm × 30 cm column, equilibrated with water). The interim product was analyzed by NMR. Subsequently, the BOS with an ADH linker attached was used for the conjugation to a OVA in the presence of carbodiimide (EDC) and purified by HPLC using HiLoad 16/600 Superdex 30 prep grade column. The formation of BOS-OVA conjugates was confirmed by size exclusion chromatography, as well as ELISA and immunoblot analysis.

### Immunization of animals

Rabbits (two animals) were immunized using a composition of BOS-TTd with the monophosphoryl lipid A (MPLA) as adjuvant. Immunization was performed subcutaneously, twice with an interval of 14 days. The sera were collected 21 days after the second immunization. Animals were housed at the animal facility of the Hirszfeld Institute of Immunology and Experimental Therapy, Wroclaw, Poland, and all the experiments were carried out according to the procedures approved by the Local Ethical Committee for Animal Experimentation. The antibodies against the BOS-TTd conjugates were tested in dot-blots and ELISA.

### Immunochemical methods

Enzyme-linked immunosorbent assay (ELISA) was performed by a modification ^24^ of the method described by Voller *et al.* ^33^ and immunoblotting as previously described ^34^. Polyclonal antibodies used for the cross-reactivity tests with *Bordetellae* exoglycans are listed in the **Supplementary Tab. 2**. Anti-BOS-TTd antibodies were analyzed in ELISA test, using BOS-OVA-conjugates as a solid-phase antigen for detection of anti-glycan specificities. The BOS-OVA conjugates were used as a solid-phase antigen to avoid carrier cross reactions. Monovalent antiserum to *Salmonella* capsular antigen Vi (Immunolab) was used as a control. A goat anti-rabbit IgG conjugated with alkaline phosphatase (Bio-Rad) as a second antibody and p-nitrophenyl phosphate and 5-bromo-4-chloro-3-indolyl phosphate/nitro blue tetrazolium for ELISA and immunoblotting, were used as the detection systems, respectively.

### FACS analysis

Bacteria, cultured as described above, were washed and suspended in PBS to an optical density of 6 x 10^8^ CFU/ml. Appropriate antisera (0.5 ml, 1/50 dilutions) were added to the bacterial suspension and incubated at 24°C overnight. Preimmune serum was employed as negative control. Bacteria were washed three times with PBS and incubated with fluorescein isothiocyanate-labeled goat anti-rabbit IgG (3 h, 24°C). Subsequently, bacteria were washed thoroughly and re-suspended in PBS containing 4% formaldehyde. An analysis of the fluorescein isothiocyanate-labeled bacteria was performed using a CytoFLEX SRT Benchtop Cell Sorter (Beckman Coulter Life Sciences, Indiana, USA). In each analysis 1 x 10^4^ bacterial cells were evaluated. Narrow-angle forward light scatter and green fluorescence emission signals were collected. Bacterial aggregates were electronically excluded on the basis of light scatter signals.

## Data availability

Data generated in this study are provided in the manuscript and its Supplementary Information.

## Supplementary Information

Supplementary Figs. 1-6 and Tables 1-4.

Reporting Summary

## Acknowledgements. Funding statement

This work was supported by the statutory funds of the Hirszfeld Institute and by National Science Centre grant 2021/05/X/NZ6/00216

## Author contributions

KU – finding of the new molecule. KU, SK, TN - concept and design of the work. KU, SK, TN - acquisition, analysis and interpretation of the data. TN - writing the original draft. KU, SK, TN - review and editing the final manuscript. All of the authors approved and contributed to the final version of the manuscript.

**The authors declare no competing interests**

