## Supplementary material for "Discovery of *Bordetella* oligosaccharide – extracellular glycan common to genus *bordetella*. Structure, immunogenicity and possible implications for host-pathogen interactions": SI: Supplementary_materials_v7.pdf

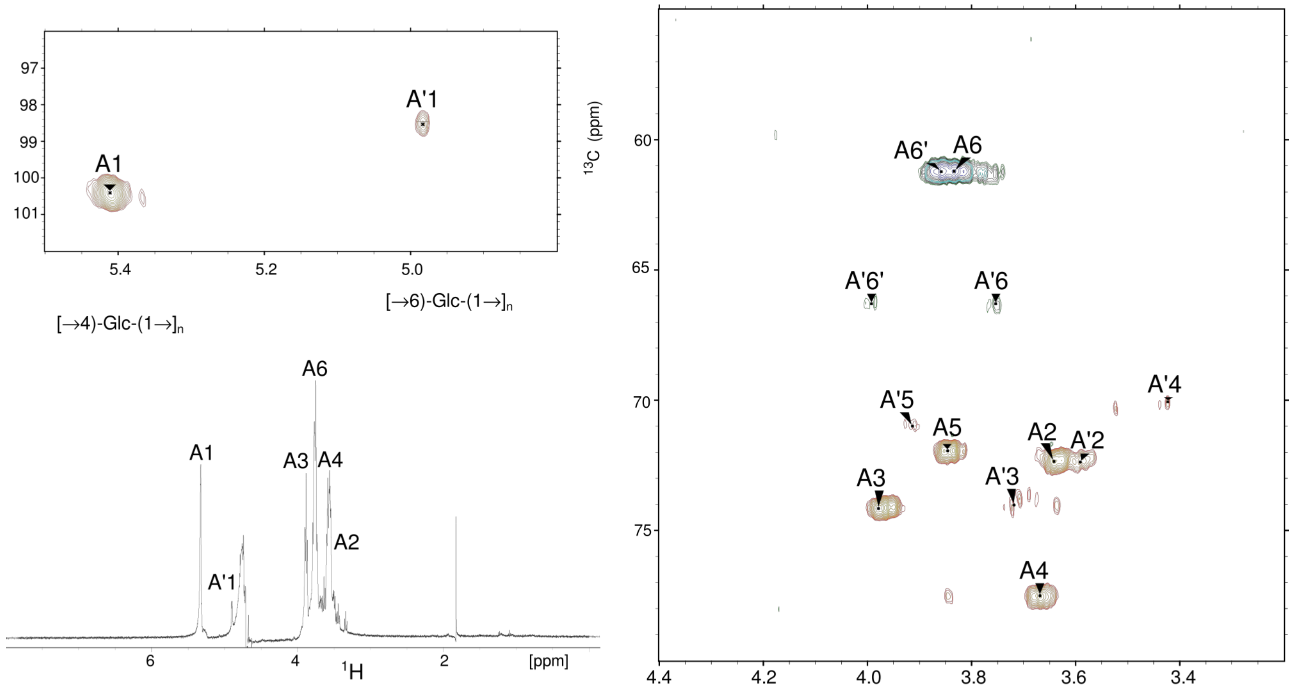


**Supplementary Fig. 1. HSQC-DEPT spectrum of the exopolysaccharide (fraction I) of *B. holmesii* ATCC 51541 isolated from the culture medium.** The spectrum was obtained for $^2H_2O$ solutions at 600 MHz and 25 °C. The capital letters refer to carbohydrate residues A and A' as defined in Tab. 1.

The NMR spectra (Supplementary Fig. 1) of fraction I indicated a glucose homopolymer. All signals and spin-systems were assigned by several two-dimensional experiments. Residue **A** with the H-1/C-1 signals at δ 5.41/100.4 ppm, $J_{H-1,H-2}$ 3.4 Hz, $J_{C-1,H-1}$ ~ 174 Hz was assigned as the 4-substituted α-D-Glc*p* on basis of the $^1H$ and $^{13}C$ chemical shifts, the large vicinal coupling constants between H-2, H-3, H-4 and H-5 protons in the sugar ring, and the characteristic downfield shift of the C-4 signal (δ 76.7 ppm). Similarly, residue **A'** with the H-1/C-1 signals at δ 4.98/98.5 ppm, $J_{H-1,H-2}$ 3.3 Hz, $J_{C-1,H-1}$ ~ 172 Hz was assigned as the 6-substituted α-D-Glc*p* on basis of the $^1H$ and $^{13}C$ chemical shifts, the large vicinal coupling constants between H-2, H-3, H-4 and H-5 protons in the sugar ring, and the characteristic downfield shift of the C-6 signal (δ 66.3 ppm). The HMBC and 2D NOESY spectra showed cross-peaks between the anomeric proton and the proton/carbon at the linkage position. The anomeric proton (H-1) of residue A at δ 5.41 ppm showed connectivities to the H-4/C-4 signals at δ 3.67/77.5 ppm. Likewise, the anomeric proton (H-1) of residue A' at δ 4.98 ppm showed connectivities to the H-6/C-6 signals at δ 3.75, 3.99/66.3 ppm. As there were no cross-peaks between the residues A and A', it was concluded that the fraction I is a mixture of two types of glucans: α-(1→4)-linked and α-(1→6)-linked homopolymers of glucose.

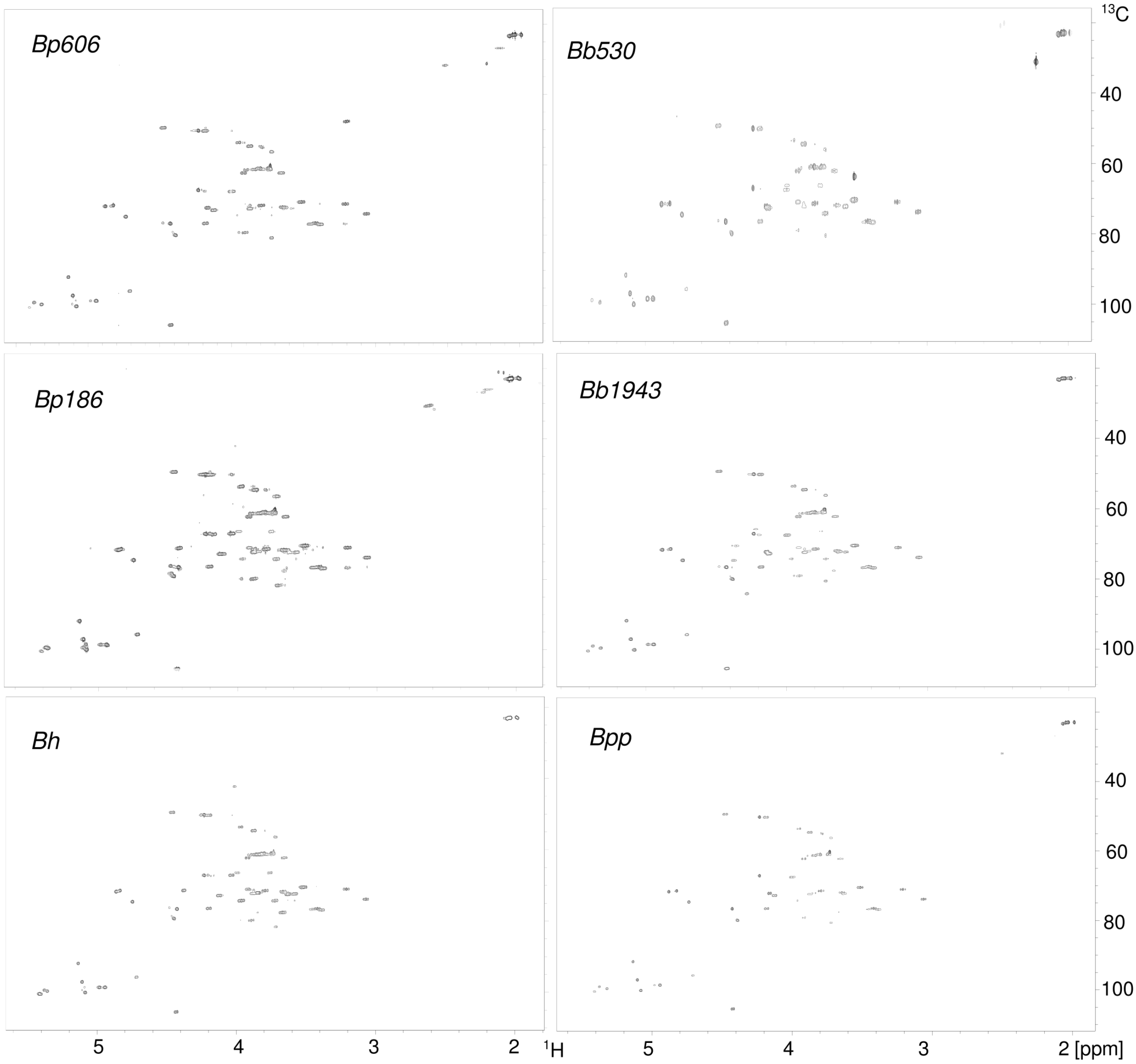


**Supplementary Fig. 2. HSQC-DEPT spectra of BOS isolated from *Bordetellae***: *B. pertussis* 606, *B. pertussis* 186, *B. holmesii ATCC 51541*, *B. bronchiseptica* 530, *B. bronchiseptica* 1943, and *B. parapertussis* 529.

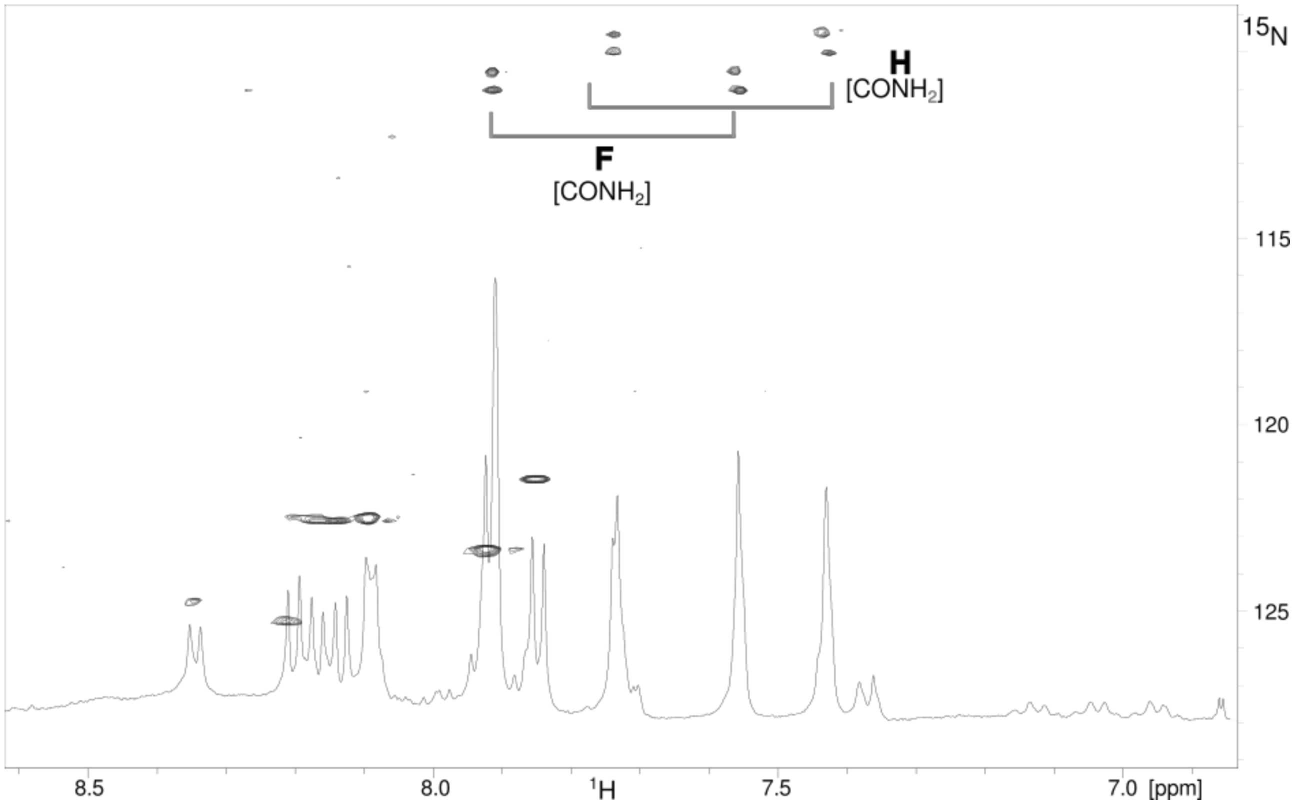


**Supplementary Fig. 3. Parts of the $^{1}H$ NMR and 2D $^{1}H,^{15}N$ HSQC spectra of exchangeable protons of amides in *Bordetellae* BOS.** The spectrum was obtained for 90% $H_2O$, prior to exchange with 10 % $^{2}H_2O$. The amide protons of residues F and H at $\delta_N$ 111 ppm and 110 ppm are indicated, respectively.

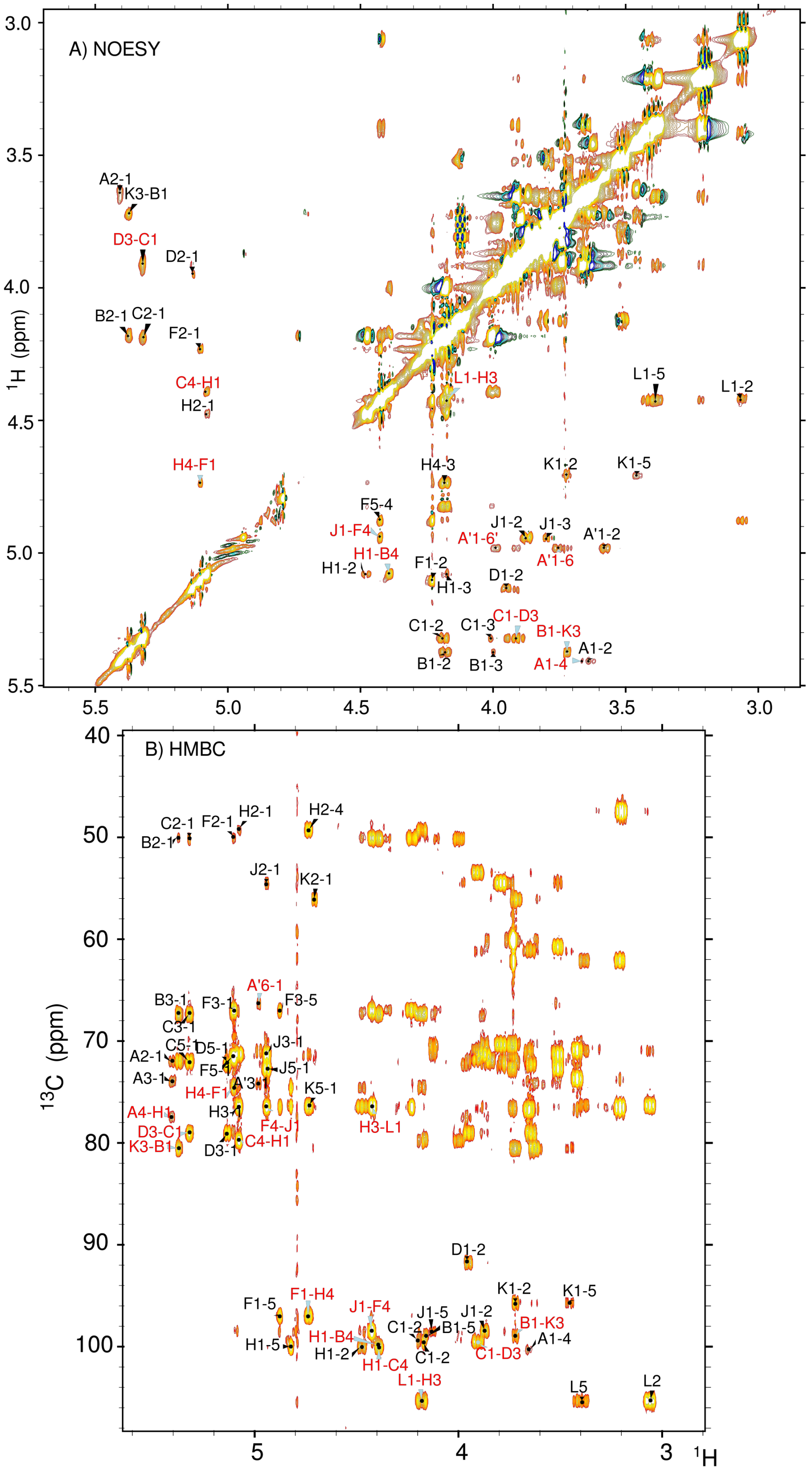


**Supplementary Fig. 4. Parts of the NOESY and HMBC spectra of the exooligosaccharide BOS of *B. pertussis*.** The spectra were obtained for $^2H_2O$ solution at 600 MHz and 25 °C. The NOE/$^3J_{C,H}$ -connectivities were recorded with 200 ms mixing-time. The delay for the evolution of long range coupling in HMBC spectra was 60 ms. The cross-peaks are labeled as shown in Fig. 3.

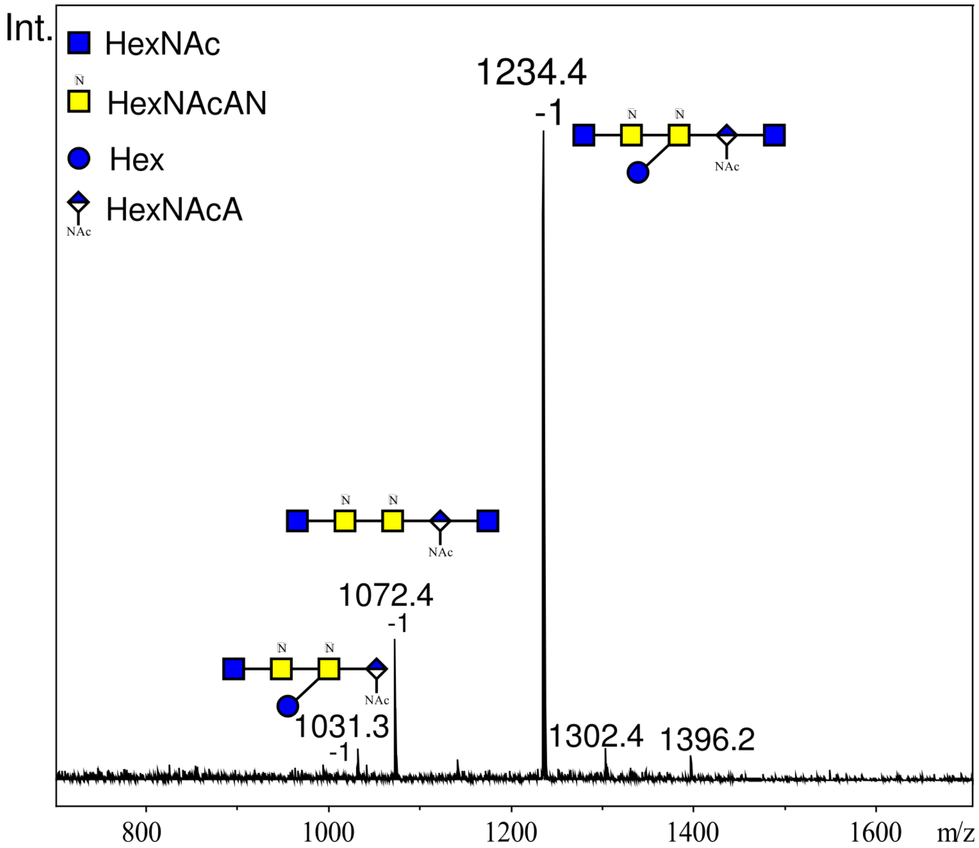


**Supplementary Fig. 5. ESI MS of the exooligosaccharide fraction III (BOS).** The spectrum was recorded in negative ion mode.

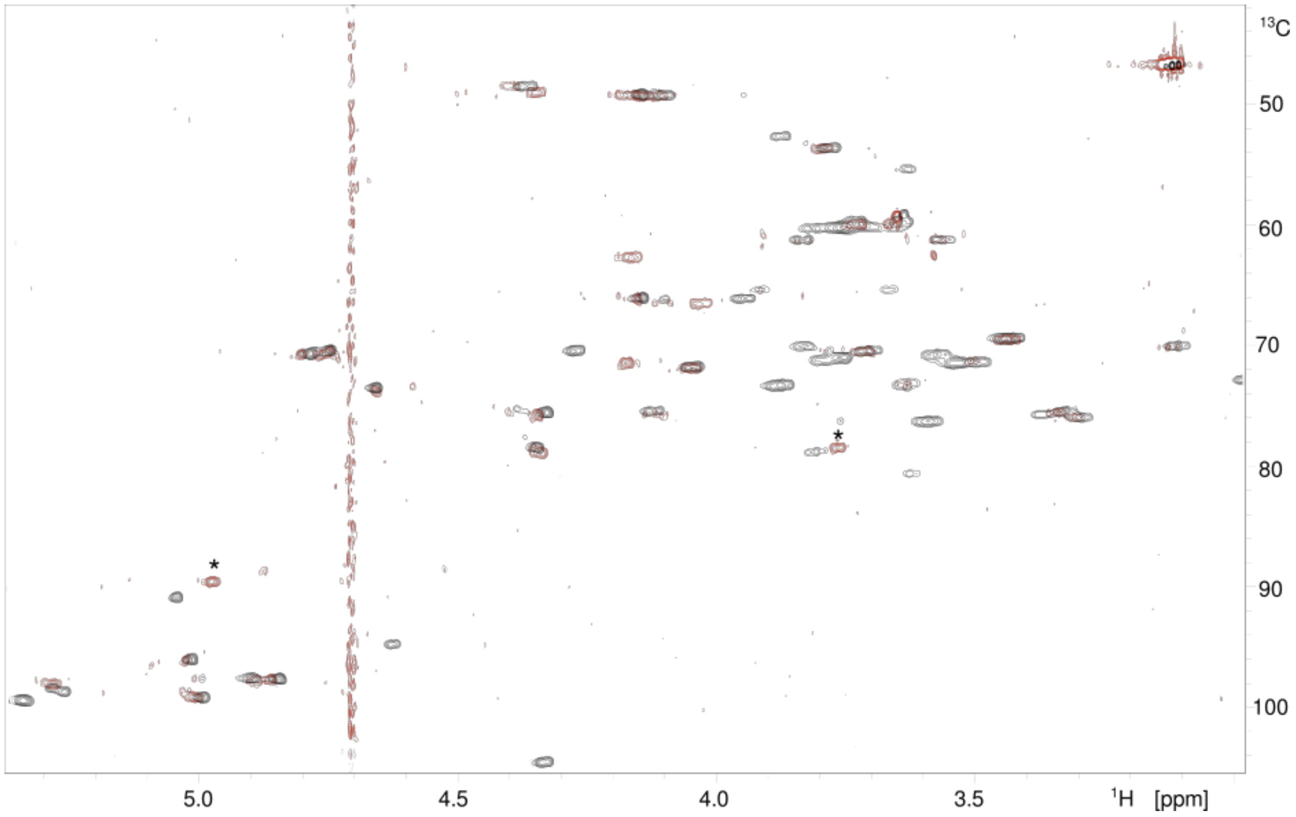


**Supplementary Fig. 6. HSQC-DEPT spectra of *B. holmesii ATCC 51541* BOS (black) and the oxidized BOS (red).** The modified signals of the oxidized residues D and K are indicated with the asterics.

**Supplementary Table 1. *Bordetellae* exoglycans quantities of the the isolated fractions**

| Bacteria (species strain) | Fraction number & the amount of material recovered [mg] | | | |
|---|---|---|---|---|
| | Fr. I | Fr. II | Fr. III | Fr. IV |
| *Bp* Tohama | 0.1 | 0.4 | 3.7 | 1.7 |
| *Bp* 186 | 0.8 | ~0.1 | 3.6 | ~0.2 |
| *Bp* 606 | ~0.1 | ~0.1 | 3.4 | 0.4 |
| *Bp* 629 | 0.9 | 0.5 | 2.5 | 4.4 |
| *Bpp* 529 | ~0.1 | ~0.1 | 3.2 | - |
| *Bh* | 1.6 | 1.2 | 1.8 | 4.8 |
| *Bb* 530 | ~0.1 | ~0.1 | 0.6 | ~0.1 |
| *Bb* 1943 | ~0.1 | 0.3 | 3.0 | 0.6 |
| *Bp* 23/68* | - | - | 0.6 | ~0.1 |
| *Bp* 9/68* | - | ~0.1 | 2.7 | 0.8 |
| *Bp* 35/68* | 2.6 | 0.8 | 9.7 | - |
| *Bp* 1310/01* | 0.8 | 0.4 | 6.1 | - |
| *Bp* 1326/62* | - | - | 0.6 | 0.2 |
| *Bp* 273/98* | ~0.1 | 0.4 | 2.8 | - |
| *Bp* 598/99* | - | - | 0.6 | ~0.1 |
| *Bp* 969/05* | - | 0.4 | 0.2 | 1 |
| *Bp* 1245/70* | - | - | 3.0 | - |
| *Bp* 2138/98* | ~0.1 | 0.4 | 1.4 | 1.1 |

(*) Asterisk denotes clinical strains, indicating a id number & a year of isolation. *Bp - B. pertussis, Bpp - B. parapertussis, Bh – B. holmesii, Bb - B. bronchiseptica.*

**Supplementary Table 2. Polyclonal antibodies used for the cross-reactivity tests with *Bordetellae* exoglycans**

| | **Serum Identifier** | **Description – antibodies against:** |
|---|---|---|
| 1 | anti–disacch-HSA | Distal disaccharide of *B. pertussis* LOS conjugated to human serum albumin as a carrier |
| 2 | anti–OS-PT | Complete core oligosaccharide of *B. pertussis* LOS conjugated to pertussis toxin as a carrier |
| 3 | anti–penta-PT | Distal pentasaccharide of *B. pertussis* LOS obtained by deamination, conjugated to to pertussis toxin as a carrier |
| 4 | anti–penta-TTd | Distal pentasaccharide of *B. pertussis* LOS obtained by deamination, conjugated to to tetanus toxoid as a carrier |
| 5 | anti–WB *Bp* 186 | Whole bacterial cells in a mixture of *B. pertussis* strains 186 |
| 6 | anti–WB *Bp* mix | Whole bacterial cells in a mixture of *B. pertussis* strains 186, 576, 606 & 629 used for vaccination (whole-cell pertussis vaccine) |
| 7 | anti-Vi | *Salmonella* antigen group Vi (capsular) for bacterial serotyping |
| 8 | anti–LPS *P.shigelloides* 78/89 | Whole bacterial cells of *P. shigelloides* 78/89 (unrelated serum control) |

**Supplementary Table 3. Bacterial strains**

| | |
|---|---|
| National Medicines Institute | *B. pertussis* 1326/62<br>*B. pertussis* 9/68<br>*B. pertussis* 23/68<br>*B. pertussis* 35/68<br>*B. pertussis* 1245/70<br>*B. pertussis* 273/98<br>*B. pertussis* 2138/ 98<br>*B. pertussis* 598 /99<br>*B. pertussis* 1310/01<br>*B. pertussis* 969/05<br>B. pertussis 186*<br>*B. pertussis* 606*<br>*B. pertussis* 629*<br>* strains used in the wP vaccine manufactured in Poland |
| Polish Collection of Microorganisms | *B. bronchiseptica* strain PCM 530 [ATCC 19395]<br>*B. bronchiseptica* strain PCM 1943 [ATCC 4617]<br>*B. parapertussis* strain PCM 529 [ATCC 15311] |
| German Collection of Microorganisms and Cell Cultures | *B. holmesii* strain DSM 13416 (ATCC 51541)<br>*B. petrii* strain DSM 12804<br>*B. hinzii* strain DSM 11333 |
| National Collection of Type Cultures | *B. pertussis* strain NCTC 13251<br>*B. pertussis* strain Tohama I |

**Supplementary Table 4. Composition of the media**

| Media | | g/L |
|---|---|---|
| **Stainer–Scholte pH 7.9** | sodium glutamate* | 10.72 |
| | L-proline | 0.24 |
| | $KHPO_4$ | 0.5 |
| | NaCl | 2.5 |
| | KCl | 0.2 |
| | $MgCl_2$ $6H_2O$ | 0.1 |
| | $CaCl_2$ | 0.02 |
| | Trishydroxymethylaminomethane | 6.1 |
| | Ryboflavine | 0.01 |
| **Charcoal agar pH 7.4** | Lab lemco powder | 10 |
| | Peptone | 10 |
| | Starch | 10 |
| | Charcoal bacteriological | 4 |
| | NaCl | 5 |
| | Nicotinic acid | 0.001 |
| | Agar | 12 |
| Growth factors | Nicotinic acid | 0.004 |
| | L-cysteine HCl $H_2O$ | 0.04 |
| | Reduced glutathion | 0.15 |
| | Ascorbic acid | 0.4 |
| | $FeSO_4$ • $7H_2O$ | 0.01 |

(*) In the metabolic labeling culture sodium glutamate was replaced by the sodium salt of $^{13}C$,$^{15}N$-labeld L-Glutamate (Silantes, Munich, Germany)
